# Multimodal longitudinal prediction of alcohol-use trajectories in adolescents and young adults: Social context, real-life social affective benefit, and fronto-striatal involvement

**DOI:** 10.64898/2026.10.28.762013

**Authors:** Oksana Berhe, Gabriela Gan, Jamila Andoh, Ali Ghadami, Markus Reichert, Ulrich W. Ebner-Priemer, Christine Heim, Falk Kiefer, Andreas Meyer-Lindenberg, Heike Tost

**Affiliations:** Department of Psychiatry and Psychotherapy, Central Institute of Mental Health, Medical Faculty Mannheim, University of Heidelberg, Mannheim, Germany; German Center for Mental Health (DZPG), Partner Site Mannheim-Heidelberg-Ulm, Mannheim, Germany; Department of Sport and Exercise Science, Faculty of Natural and Life Sciences, University Salzburg, Salzburg, Austria; Mental mHealth Lab, Department of Applied Psychology, Institute of Sports and Sports Science, Karlsruhe Institute of Technology, Karlsruhe, Germany; Charité-Universitätsmedizin Berlin, Corporate Member of Freie Universität Berlin and Humboldt-Universität zu Berlin, Institute of Medical Psychology, Berlin, Germany; German Center for Mental Health (DZPG), Partner Site Berlin-Potsdam, Germany; Charité-Universitätsmedizin Berlin, Corporate Member of Freie Universität Berlin and Humboldt-Universität zu Berlin, NeuroCure Cluster of Excellence, Berlin, Germany; Department of Addictive Behavior and Addiction Medicine, Central Institute of Mental Health, Medical Faculty Mannheim, University of Heidelberg, Mannheim, Germany

**Author notes:** **Corresponding Author** Dr. Oksana Berhe, Central Institute of Mental Health, Department of Psychiatry and Psychotherapy, Square J5, 68159 Mannheim, Germany. These authors contributed equally (Andreas Meyer-Lindenberg, Heike Tost).

**Keywords:** Adolescent and young adult alcohol use, social-affective benefit, ecological momentary assessment, reward anticipation, ventral striatum, longitudinal study

## Abstract

Escalating alcohol use during adolescence and young adulthood confers substantial risk for later substance misuse and adverse mental health outcomes, yet the factors that determine early drinking trajectories remain poorly understood. Here, we examined alcohol use across social, affective, and neural domains in a cohort of 378 community-based young drinkers using a combination of longitudinal phenotyping, ecological momentary assessment (EMA) and functional neuroimaging. Across different levels of analysis, higher baseline alcohol consumption (standard drinks per week) was associated with immediate substance-use social context, greater social-affective benefit in daily life, and heightened engagement of the fronto-striatal reward circuitry. In a one-year longitudinal follow-up, individual social-affective benefit and substance-related social environmental factors showed the strongest predictive value for drinking trajectories in a follow-up of 219 individuals after one year. Our findings identify key social and neural reward-related determinants of youth drinking behavior, informing the development of evidence-based risk detection and prevention measures.

## Introduction

The transition from adolescence to early adulthood is marked by profound life changes – leaving parental home, establishing new peer networks, and entering less supervised social environments where alcohol is widely available and socially accepted. While increasing social engagement during this period of life is a process of developmental adaptation, it also coincides with the potential for a pronounced escalation in alcohol use, which confers risk for alcohol abuse and adverse mental health outcomes later in life^1, 2^. Despite its high public health relevance, the factors that shape youth drinking trajectories remain poorly understood. Youth drinking has declined across most high-income countries from the mid-2000s onwards^3^. Against this backdrop, identifying modifiable social and affective determinants of drinking trajectories in community-based young drinkers is crucial to understand the factors that may lead individuals to deviate from the normative trend.

A defining characteristic of youth drinking is its social nature. During this life stage, alcohol consumption typically occurs in social settings with peers, such as parties, nightlife venues, or gatherings with friends^4^. Here, the initiation and continuation of alcohol use are strongly influenced by social motives, including the behavior of role models and peer expectations^5, 6^. Notably, studies have consistently shown that being in the company of others is associated with greater positive affect and reduced negative affect, yielding measurable emotional benefits in everyday life^7, 8^. Converging evidence further suggests that alcohol use and social experience interact within this affective landscape. Individuals frequently report drinking to enhance sociability and interpersonal bonding^9^, while social contexts amplify the subjective pleasurable effects of alcohol^10, 11^. Recent social-cognitive models conceptualize alcohol use disorder risk as emerging from repeated coupling of drinking with socially rewarding contexts, which progressively enhances the incentive salience of alcohol-related cues and situations. Extending this account, studies further show that individuals who experience stronger rewarding effects from alcohol, particularly under socially facilitating conditions, are at a higher risk for problematic alcohol use later in life^11^. Our multimodal design allows us to test key elements of this framework in a non-clinical, developmental sample.

At the neural level, the ventral striatum (VS) is central to reward-related processes. The VS is a core hub of the mesolimbic reward system that is involved in the processing of incentive motivation and reward expectancy^12, 13^. Neuroimaging studies have consistently demonstrated altered striatal responses during reward processing in alcohol use disorder and other addictions^14-16^. The VS is also linked to alcohol cue reactivity and the initiation, escalation, and maintenance of drinking behavior^14, 16-18^. Importantly, the social context itself modulates this system: peer presence enhances striatal reward responses, linking social experience to neural reward processing^19^. Furthermore, the interaction between the VS and prefrontal cortical regions is essential for integrating motivational value of cues with decision-making and behavioral regulation. Accordingly, altered functional connectivity within this circuitry has been linked to alcohol abuse and addiction^20, 21^.

Taken together, the literature consistently shows that alcohol use is embedded within a broader matrix of incentive processes spanning social environment, everyday affective experience, and neural reward processing. While multimodal and longitudinal approaches are increasingly being adopted in youth alcohol research^22-24^, the evidence remains fragmented and often limited to unimodal and/or cross-sectional studies. Specifically, epidemiological studies typically characterize drinking through demographic or environmental risk factors^25, 26^, whereas neuroimaging studies focus on reward-related neural alterations in laboratory paradigms^21, 27^. Ambulatory assessment studies, in turn, capture individual affective experience in real-world settings but seldom integrate neural function or trajectory dynamics^10^. Additionally, research on youth is limited and primarily focuses on clinical or high-risk populations (e.g., alcohol use disorder (AUD), heavy drinkers, or familiar risk)^21^ with rare attention paid to healthy community-based young drinkers^24^.

To account for these limitations, this study applied a multimodal, longitudinal, and ecologically grounded neurobehavioral approach to a community-based cohort of healthy adolescents and young adults with occasional alcohol use and no prior psychiatric disorders to examine reward-related and social determinants of drinking before the emergence of clinically significant impairment. Towards this aim, we focused on a group of individuals who consume alcohol, but in low to moderate amounts (mean ± standard deviation (SD): 2.65 ± 2.76 standard drinks per week), i.e., less than the typical high-risk drinkers described in the literature. Of note, none of the participants had any relevant current or former psychiatric symptoms, diagnoses, or treatments^28^. Focusing on this population allowed us to capture early, potentially modifiable risk processes that are often obscured in treatment-seeking or high-risk samples. Using a multimodal approach, we integrated questionnaire-based, real-life, and neuroimaging measures, to comprehensively characterize the relationship of these measures to current alcohol use and assess their predictive value for changes in drinking behavior from baseline to follow-up. At baseline, we first identified individual and social-contextual factors associated with drinking behavior and examined their relationship with social-affective experience in daily life. We then examined fronto-striatal activity and connectivity during reward anticipation using functional magnetic resonance imaging (fMRI) and a validated reward-processing task^29, 30^. Finally, we examined how the identified multimodal markers related to prospective changes in alcohol use over the course of roughly one year (mean ± standard deviation (SD) of follow-up measurements: 13.25 ± 8.4 months). We hypothesized that higher alcohol use would be associated with proximal social and substance-related environmental factors, a higher social-affective reinforcement in daily life, and an increased engagement of the fronto-striatal reward circuitry.

## Results

### Psychological features

#### Baseline correlates

Quantity of consumed alcohol was significantly higher among older individuals and males (all p values < 0.001). Multiple linear regression analysis revealed that risky substance-use environments, i.e. peer environments with a higher propensity for substance use (ESPAD questionnaire^31^; ß = 0.344, p < 0.001) and greater individual polysubstance use (ESPAD, ß = 0.242, p < 0.001) were the strongest predictors of current alcohol consumption after correcting for age and sex (Table 1, Fig.1). Additionally, increased social network size (Social Network Index, SNI^32^, ß = 0.176, p = 0.004) and elevated daily stress levels (daily stress inventory, ABF^33^, ß = 0.140, p < 0.007) were identified as significant predictors. These findings remained significant after adjusting for legal alcohol access. In contrast, several well-established factors, such as socioeconomic status (SES) and familial psychiatric history, showed no significant association with consumption quantity in our sample (all p values > 0.62).

**Table 1.**
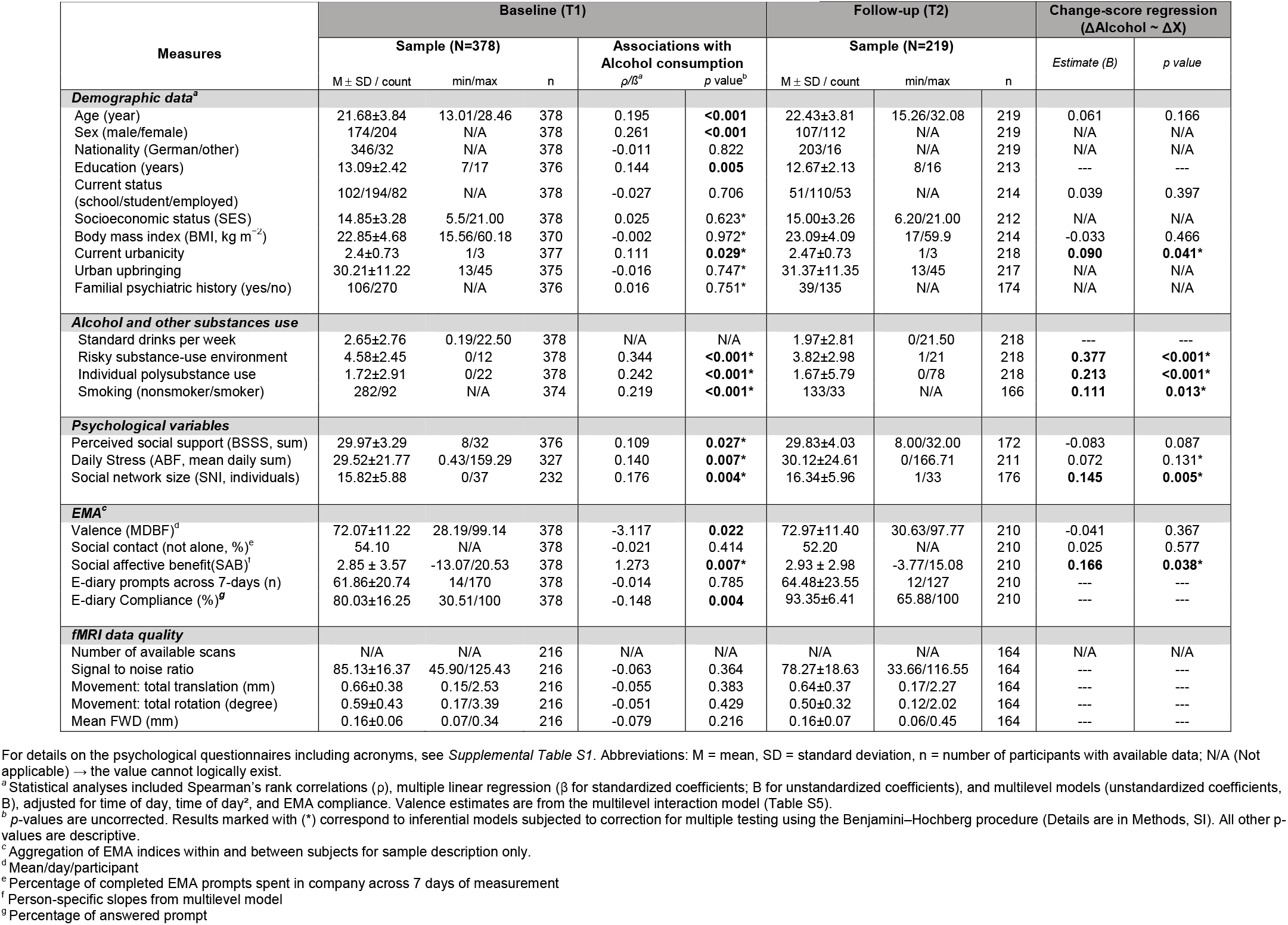
Sample Characteristics and Change-Score Regression Results.

| Measures | Baseline (T1) |  |  |  |  | Follow-up (T2) |  |  | Change-score regression (ΔAlcohol ~ ΔX) |  |
|---|---|---|---|---|---|---|---|---|---|---|
|  | Sample (N=378) |  |  | Associations with Alcohol consumption |  | Sample (N=219) |  |  | *Estimate (B)* | *p value* |
|  | M ± SD / count | min/max | n | *ρ/ß*<sup>a</sup> | p value<sup>b</sup> | M ± SD / count | min/max | n |  |  |
| ***Demographic data<sup>a</sup>*** |  |  |  |  |  |  |  |  |  |  |
| Age (year) | 21.68±3.84 | 13.01/28.46 | 378 | 0.195 | **<0.001** | 22.43±3.81 | 15.26/32.08 | 219 | 0.061 | 0.166 |
| Sex (male/female) | 174/204 | N/A | 378 | 0.261 | **<0.001** | 107/112 | N/A | 219 | N/A | N/A |
| Nationality (German/other) | 346/32 | N/A | 378 | -0.011 | 0.822 | 203/16 | N/A | 219 | N/A | N/A |
| Education (years) | 13.09±2.42 | 7/17 | 376 | 0.144 | **0.005** | 12.67±2.13 | 8/16 | 213 | --- | --- |
| Current status (school/student/employed) | 102/194/82 | N/A | 378 | -0.027 | 0.706 | 51/110/53 | N/A | 214 | 0.039 | 0.397 |
| Socioeconomic status (SES) | 14.85±3.28 | 5.5/21.00 | 378 | 0.025 | 0.623* | 15.00±3.26 | 6.20/21.00 | 212 | N/A | N/A |
| Body mass index (BMI, kg m<sup>−2</sup>) | 22.85±4.68 | 15.56/60.18 | 370 | -0.002 | 0.972* | 23.09±4.09 | 17/59.9 | 214 | -0.033 | 0.466 |
| Current urbanicity | 2.4±0.73 | 1/3 | 377 | 0.111 | **0.029\*** | 2.47±0.73 | 1/3 | 218 | **0.090** | **0.041\*** |
| Urban upbringing | 30.21±11.22 | 13/45 | 375 | -0.016 | 0.747* | 31.37±11.35 | 13/45 | 217 | N/A | N/A |
| Familial psychiatric history (yes/no) | 106/270 | N/A | 376 | 0.016 | 0.751* | 39/135 | N/A | 174 | N/A | N/A |
| ***Alcohol and other substances use*** |  |  |  |  |  |  |  |  |  |  |
| Standard drinks per week | 2.65±2.76 | 0.19/22.50 | 378 | N/A | N/A | 1.97±2.81 | 0/21.50 | 218 | --- | --- |
| Risky substance-use environment | 4.58±2.45 | 0/12 | 378 | 0.344 | **<0.001\*** | 3.82±2.98 | 1/21 | 218 | **0.377** | **<0.001\*** |
| Individual polysubstance use | 1.72±2.91 | 0/22 | 378 | 0.242 | **<0.001\*** | 1.67±5.79 | 0/78 | 218 | **0.213** | **<0.001\*** |
| Smoking (nonsmoker/smoker) | 282/92 | N/A | 374 | 0.219 | **<0.001\*** | 133/33 | N/A | 166 | **0.111** | **0.013\*** |
| ***Psychological variables*** |  |  |  |  |  |  |  |  |  |  |
| Perceived social support (BSSS, sum) | 29.97±3.29 | 8/32 | 376 | 0.109 | **0.027\*** | 29.83±4.03 | 8.00/32.00 | 172 | -0.083 | 0.087 |
| Daily Stress (ABF, mean daily sum) | 29.52±21.77 | 0.43/159.29 | 327 | 0.140 | **0.007\*** | 30.12±24.61 | 0/166.71 | 211 | 0.072 | 0.131* |
| Social network size (SNI, individuals) | 15.82±5.88 | 0/37 | 232 | 0.176 | **0.004\*** | 16.34±5.96 | 1/33 | 176 | **0.145** | **0.005\*** |
| ***EMA<sup>c</sup>*** |  |  |  |  |  |  |  |  |  |  |
| Valence (MDBF)<sup>d</sup> | 72.07±11.22 | 28.19/99.14 | 378 | -3.117 | **0.022** | 72.97±11.40 | 30.63/97.77 | 210 | -0.041 | 0.367 |
| Social contact (not alone, %)<sup>e</sup> | 54.10 | N/A | 378 | -0.021 | 0.414 | 52.20 | N/A | 210 | 0.025 | 0.577 |
| Social affective benefit(SAB)<sup>f</sup> | 2.85 ± 3.57 | -13.07/20.53 | 378 | 1.273 | **0.007\*** | 2.93 ± 2.98 | -3.77/15.08 | 210 | **0.166** | **0.038\*** |
| E-diary prompts across 7-days (n) | 61.86±20.74 | 14/170 | 378 | -0.014 | 0.785 | 64.48±23.55 | 12/127 | 210 | --- | --- |
| E-diary Compliance (%)<sup>***g***</sup> | 80.03±16.25 | 30.51/100 | 378 | -0.148 | **0.004** | 93.35±6.41 | 65.88/100 | 210 | --- | --- |
| ***fMRI data quality*** |  |  |  |  |  |  |  |  |  |  |
| Number of available scans | N/A | N/A | 216 | N/A | N/A | N/A | N/A | 164 | N/A | N/A |
| Signal to noise ratio | 85.13±16.37 | 45.90/125.43 | 216 | -0.063 | 0.364 | 78.27±18.63 | 33.66/116.55 | 164 | --- | --- |
| Movement: total translation (mm) | 0.66±0.38 | 0.15/2.53 | 216 | -0.055 | 0.383 | 0.64±0.37 | 0.17/2.27 | 164 | --- | --- |
| Movement: total rotation (degree) | 0.59±0.43 | 0.17/3.39 | 216 | -0.051 | 0.429 | 0.50±0.32 | 0.12/2.02 | 164 | --- | --- |
| Mean FWD (mm) | 0.16±0.06 | 0.07/0.34 | 216 | -0.079 | 0.216 | 0.16±0.07 | 0.06/0.45 | 164 | --- | --- |

For details on the psychological questionnaires including acronyms, see *Supplemental Table S1*. Abbreviations: M = mean, SD = standard deviation, n = number of participants with available data; N/A (Not applicable) → the value cannot logically exist.
<sup>*a*</sup> Statistical analyses included Spearman's rank correlations (ρ), multiple linear regression (β for standardized coefficients; B for unstandardized coefficients), and multilevel models (unstandardized coefficients, B), adjusted for time of day, time of day², and EMA compliance. Valence estimates are from the multilevel interaction model (Table S5).
<sup>*b*</sup> *p*-values are uncorrected. Results marked with (\*) correspond to inferential models subjected to correction for multiple testing using the Benjamini–Hochberg procedure (Details are in Methods, SI). All other p-values are descriptive.
<sup>*c*</sup> Aggregation of EMA indices within and between subjects for sample description only.
<sup>d</sup> Mean/day/participant
<sup>e</sup> Percentage of completed EMA prompts spent in company across 7 days of measurement
<sup>f</sup> Person-specific slopes from multilevel model
<sup>g</sup> Percentage of answered prompt

#### Longitudinal prediction

Alcohol consumption quantity (standard drinks/week, untransformed) significantly decreased from baseline to follow-up (*t*_(217)_ = 3.88, p <0.001; Fig. 1A). The effect remained significant after adjustment for age, sex and legal alcohol access in a linear mixed model (F_(1,217)_ = 12.53, p <0.001). Within-person increase in risky substance-use environment, individual polysubstance use and social network size were significantly associated with increase in alcohol use over time (β = 0.377, β = 0.213, β = 0.145, all p values ≤ 0.005; Table 1, Fig. 1B-D). Changes in the other examined psychological factors did not relate to prospective changes in alcohol use over time (all p-values ≥.087; Table 1, Table S4).

**Figure 1.**
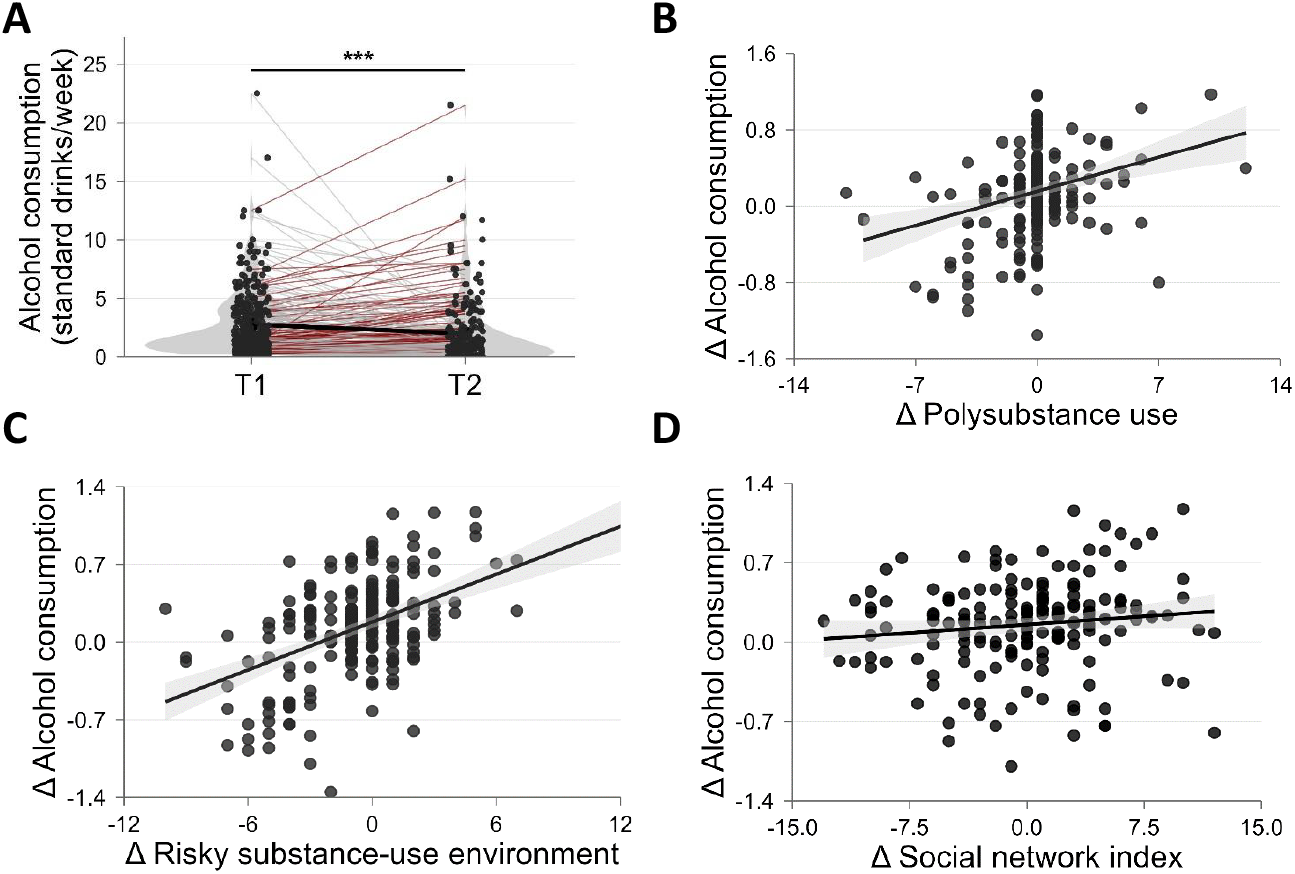
Predictors of alcohol consumption at baseline (T1) and of change in alcohol use over time (Δ T2–T1). **A**. Distribution of alcohol consumption on the original scale (standard drinks/week) at T1 and T2 shown as violin plots with individual trajectories (red = increase; grey = decrease/stable) and the paired mean trajectory (black), demonstrating a significant overall decrease (***, P < 0.001). **B–D**. Scatterplots of significant longitudinal predictors showing associations between changes in predictors and changes in log-transformed alcohol consumption (Δ T2–T1), with fitted regression lines (solid lines) and 95% confidence bands (shaded areas).

### Daily-life assessments

#### Baseline correlates

‘Social affective benefit’ (SAB) refers to the phenomenon of individuals experiencing higher affective valence in their daily lives when they are in company compared to being alone^7^. In our study, participants with higher alcohol consumption showed significantly increased SAB in daily life (ß = 1.27, p = 0.007, Fig. 2A, Table S5). This interaction occurred alongside a significant negative main effect of drinking quantity on affective valence in daily life (ß = -3.12, p = 0.022). Both results were robust to adjustment for legal alcohol access. Statistical robustness checks support the observed association (Table 1, Fig. S3).

**Figure 2.**
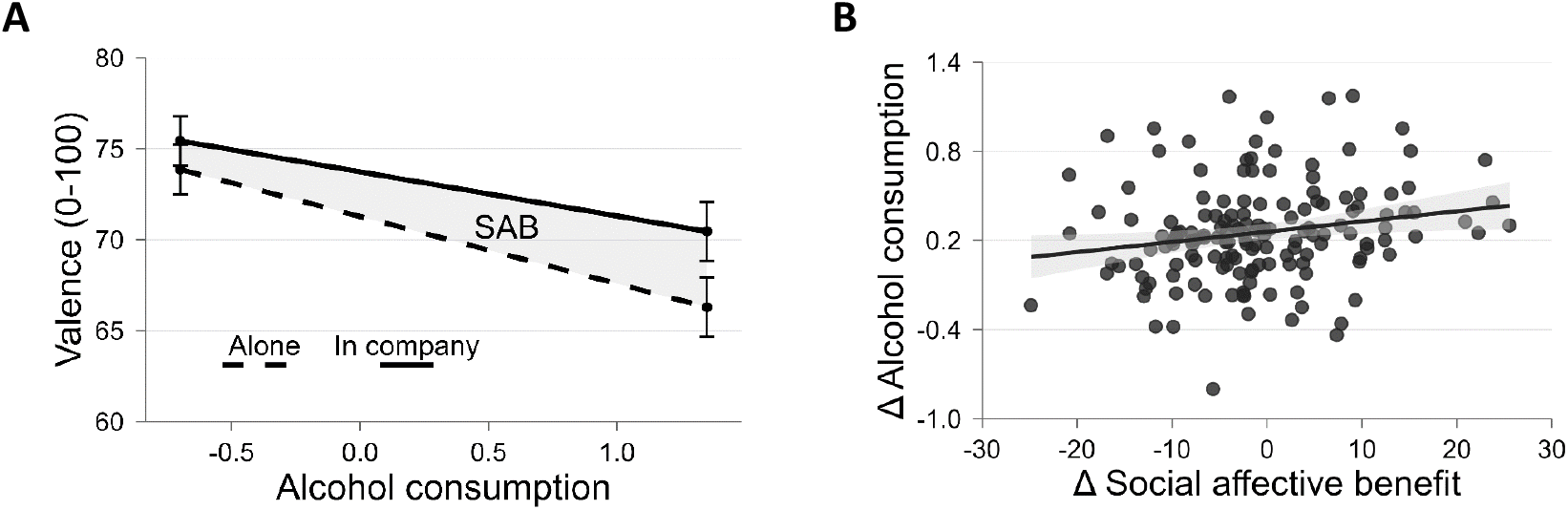
**A**. Higher log alcohol consumption (x-axis) relates to greater social affective benefit (SAB) at baseline (T1), illustrated by the shaded area between the solid (in company) and dashed (alone) lines. Regression lines are derived from multilevel interaction analyses (outcome: affective valence; predictor: real-life social contact; moderator: log alcohol consumption). Error bars represent the standard error (SE) of the estimated mean valence scores. **B**. Changes in social affective benefit predict changes in log alcohol consumption over time (Δ T2–T1), with fitted regression line and 95% confidence band (shaded areas).

#### Longitudinal prediction

Within-subject increases in SAB were significantly related to increases in alcohol use from T1 to T2 (ß = 0.166, p = 0.038; Table 1, Fig. 2B). This suggests that the more individuals were able to increase their affective state through social contact in daily life over time, the more they prospectively increased their alcohol consumption, or vice versa.

### Neural reward anticipation

#### Baseline correlates

Higher alcohol consumption at baseline was associated with increased VS activity (t = 2.84, p_FWE_ = 0.044, Fig. 3A) and stronger fronto-striatal PPI connectivity (VS-dlPFC) during reward anticipation (whole-brain corrected: t = 4.62, p_FWE_ = 0.044, Fig. 3B). No significant moderation by age or sex was detected (all p_FWE_ ≥ 0.485; Supplementary Results).

**Figure 3.**
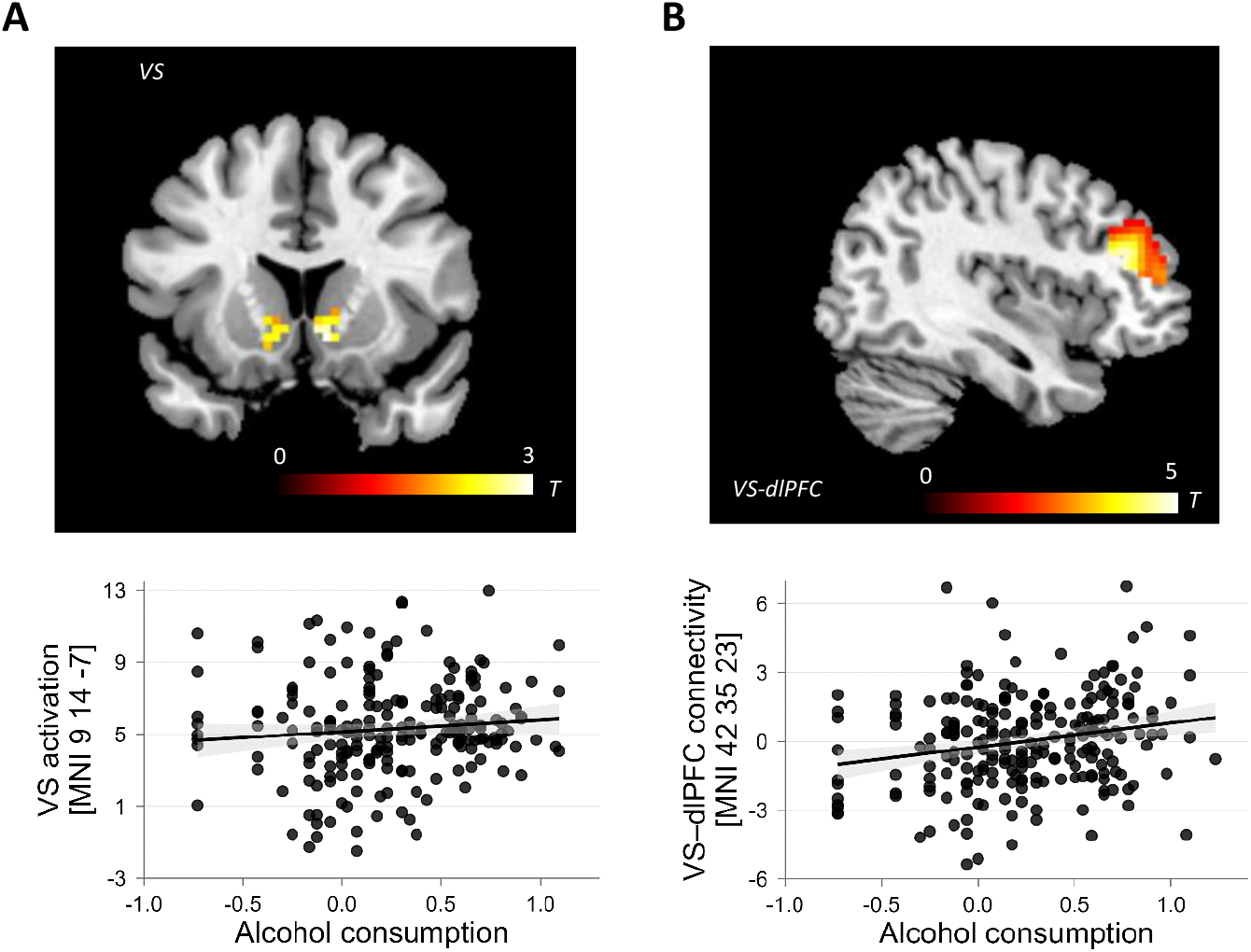
Alcohol consumption, VS activation and fronto-striatal functional connectivity at baseline. **A. Upper panel:** Brain map showing a positive association between log alcohol consumption and ventral striatal activation during reward anticipation (MNI peak voxel coordinates: 9, 14, −7, t = 2.84, pFWE = 0.044, region-of-interest corrected). **Lower panel:** Scatterplot illustrating the relationship between log alcohol consumption and VS activation at the peak voxel, with fitted regression line and 95% confidence band (shaded areas). **B. Upper panel:** Brain map showing a positive association between log alcohol consumption and VS– dlPFC functional connectivity during reward anticipation, derived from psychophysiological interaction (PPI) analysis (MNI peak voxel coordinates: 42, 35, 23, t = 4.62, pFWE = 0.044, whole-brain corrected). **Lower panel:** Scatterplot depicting the association between log alcohol consumption and VS–dlPFC functional connectivity at the peak voxel, with fitted regression line and 95% confidence band (shaded areas). For visualization purposes, statistical maps are displayed at p < .005 (uncorrected).

#### Longitudinal prediction

Longitudinal analyses revealed no significant relationships between changes in alcohol consumption and changes in VS activity or connectivity over time (all p values >.269). Complementary test–retest analyses indicated moderate reliability of VS activation (ICC = .50–.51) and VS–dlPFC connectivity (ICC = .43–.46), consistent with previous findings^34^ (Supplementary Results).

## Discussion

In this multimodal study, we examined the predictors of current and prospective alcohol use among healthy young community members who are in a critical time window for the potential development of problematic alcohol use later in life^1, 2^. Three central findings emerged across levels of analysis. First, alcohol use was strongly associated with proximal social environmental factors: Beyond the own substance-related behavior, only the substance use behavior of peers and the social network size as well as their change across time predicted both current and prospective drinking. Second, greater social-affective benefit in individuals with higher alcohol use, both at baseline and as a change over time, indicates that social interaction may be a strong incentive for consuming alcohol in real life. Third, at the neural level, higher drinking levels were significantly associated with increased VS activity and stronger VS-dlPFC connectivity during reward anticipation, but the change in these markers over time did not predict prospective changes in drinking behavior.

On a psychological level, our findings corroborate a well-documented observation in the literature: The drinking behavior of adolescents and young adults is strongly influenced by their immediate social environment^6, 35^. Here, we extend the existing literature by showing that, in this age range, the prospective increase of alcohol use goes along with the escalation of substance-related behaviors of peers and a growing size of their own social network^6, 25, 36^. This is consistent with the idea that, as young people become more independent from their family, they may become more susceptible to potentially harmful influences by peers^25, 35^. The observed association may reflect both socialization and selection: young people may adopt their peers’ drinking behavior, select drinking-oriented networks, or share common contextual and dispositional liabilities. This aligns with longitudinal network studies showing that peer drinking predicts subsequent increases in individual alcohol use^25, 35, 36^, and extends this work by demonstrating parallel within-person change in peer exposure and drinking over time. At the same time, expanding social networks may increase opportunities for exposure to drinking behavior within these contexts^6, 25, 36^.

Notably, other more time-stable demographic risk factors for alcohol abuse, in particular SES and familial psychiatric history, showed no association with current drinking behavior in our sample^26, 37^. While these factors remain important at the population level and in clinical samples, our results highlight the importance of proximal social affective processes: peer social exposures may be more strongly aligned with the own drinking behavior^35^, and might dominate the dynamic social reinforcement context in which the drinking behavior of others is shaped^4^.

The EMA findings of this study link social environmental influences to alcohol use. They revealed that adolescents and young adults who drink more showed stronger affective benefits when they were socially engaged^7, 8^. This pattern is consistent with a social-reinforcement mechanism, although the adopted observational design cannot determine whether social reward increases drinking, drinking increases the perceived value of social contact, or both processes are driven by a third factor. Moreover, within-person increases in social-affective benefit co-varied with increases in alcohol use over time. Prior research has consistently documented that social context amplifies the subjective rewarding effects of alcohol^10, 38^ and increases drinking likelihood^9^. Our findings extend this literature by demonstrating that individual differences in drinking trajectories are associated with the affective value attributed to everyday social experiences themselves. Specifically, individuals with prospectively increasing alcohol consumption appear to become more responsive to the emotional consequences of social engagement, experiencing a larger affective contrast between social and non-social contexts. These findings suggest that the drinking of adolescents and young adults becomes increasingly embedded within socially rewarding experiences over time^39^. When situated within substance-related peer environments, social experiences may reinforce continued alcohol use and contribute to escalation over time^4^. This interpretation is consistent with theoretical models proposing that repeated alcohol use strengthens the motivational significance of reward-associated cues and contexts^4, 18, 39^. Importantly, our data also show that individuals with relatively higher alcohol consumption showed lower momentary valence in daily life. Taken together, we conclude from these data that the established link between alcohol consumption and reduced well-being^40^ extends to the age range of adolescents and young adults and to real-life contexts. Moreover, we suggest that the affective advantage from real-life social interactions does not translate into more favorable emotional functioning. The cohort-wide reduction in drinking provides an ecologically valid context for interpreting our prospective associations: Factors that co-vary with increasing drinking, despite the general downward trend, may be particularly relevant for identifying youth who deviate from the typical prospective pattern. From a prevention perspective, ambulatory monitoring of social-affective benefit could help identify youths whose drinking becomes increasingly coupled to socially rewarding experiences, enabling timely, context-sensitive interventions before clinically significant problems emerge.

Our neuroimaging findings highlight a plausible biological framework for contextualizing the psychological and daily-life correlates of alcohol consumption among adolescents and young adults. Higher alcohol consumption was associated with greater striatal activity during social and monetary reward anticipation. Prior research has consistently linked altered striatal responses to alcohol consumption behavior, from risky drinking to alcohol use disorder^14-16, 41^. Further, the heightened responsivity of the neural reward system is a characteristic phenotype in youth^42^ that predicts subsequent increases in drinking^16^. Our findings extend this continuum model by showing that individual differences of ventral striatal responsivity explain differences in drinking behavior among healthy young drinkers with overall low-to-moderate consumption. This is consistent with the notion that reward sensitivity is a transdiagnostic and trait-like dimension relevant across the full severity range. Beyond regional activation, drinking was also related to stronger VS-dlPFC functional connectivity, indicating increased integration between motivational and regulatory systems during reward anticipation. In AUD, VS hyperactivity is frequently accompanied by diminished VS-dlPFC coupling, consistent with impaired fronto-striatal integration^20, 43, 44^. In healthy adults, increased fronto-striatal connectivity has been interpreted as successful regulation of reward-related responses^45^. In adolescents, however, stronger connectivity between reward and control networks has been proposed to reflect compensatory recruitment in the context of substance use^46^. Accordingly, the stronger VS-dlPFC coupling alongside increased VS activation observed in our youth sample may reflect successful recruitment of regulatory circuitry in response to heightened reward signaling, potentially indicating greater regulatory demand ^46^. Notably, prospective changes in these neural markers did not relate to changes in the alcohol consumption pattern of individuals. This may reflect the relatively modest change in alcohol consumption during the follow-up period, which may have been insufficient to induce detectable alterations in reward-related neural function in this sample^27, 47^. Alternatively, these neural markers may represent more stable, trait-like characteristics that make them more suitable for baseline prediction than longitudinal projection in this age- and/or drinking severity range. This interpretation is indirectly corroborated by the relative stability of neuroimaging phenotypes over time in our supplementary test–retest reliability analysis (Supplementary Results, Section 3.3), as well as in previous reliability studies employing the same^29^ or similar^34^ fMRI paradigms. It is important to note that the observed activation and connectivity findings should be interpreted as neural correlates of drinking quantity, rather than as evidence that hyperresponsivity of the reward circuitry causes alcohol escalation. At the same time, the robustness of these effects across age and sex strengthens their potential relevance for transdiagnostic models of reward processing and early alcohol involvement, and supports their consideration in broadly applicable prevention frameworks.

The present findings should be interpreted in light of several limitations. First, the quantity of consumed alcohol was retrospectively assessed as standard drinks per week rather than momentary intake^31^, precluding linkage of specific drinking episodes to real-time social affective experience. Second, longitudinal associations were examined using change-score regression, which identifies factors that vary in tandem with alcohol use over time but does not establish the direction of these associations^48^. Third, the relatively low and positively skewed alcohol consumption may have attenuated effect estimates and limited statistical power. Finally, the non-clinical sample is both a strength and a limitation of this study: It enabled the identification of predictors of drinking behavior in community individuals at risk but limits the generalizability of findings to clinical populations^21^. In addition, the cohort lacked comprehensive measures of alcohol-related consequences, craving, drinking motives, binge-pattern drinking, tolerance, and family history of AUD. The findings therefore pertain to early alcohol-use trajectories and social reinforcement, rather than clinically defined addiction liability. Future studies combining momentary assessments of alcohol consumption with multi-wave longitudinal designs across a broader spectrum of drinking behavior could help address these limitations.

## Conclusions

Our multimodal longitudinal study shows that alcohol consumption in healthy adolescents and young adults is consistently associated with social and reward-related determinants across behavioral, ecological, and neural levels of analysis. In a psychiatrically healthy community cohort, individual differences in drinking quantity were associated with proximal peer substance-use exposure, the affective value of everyday social contact, and fronto-striatal reward-anticipation circuitry. Prospective associations were strongest for changes in social context and social-affective experience, whereas neural measures showed cross-sectional but not longitudinal associations. The findings support the view that early processes of social reinforcement in real-life and reward sensitivity may contribute to the development of problematic alcohol use trajectories in young drinkers. By integrating these complementary levels of evidence, the present study provides a coherent multi-level characterization of risk factors for current alcohol use in young drinkers and may inform evidence-based approaches to early risk detection and prevention.

## Methods

### Study population and design

Data were drawn from two waves of the PEZ Study, a longitudinal cohort of adolescents and young adults conducted at the Central Institute of Mental Health in Mannheim, Germany (2014 -2020). At baseline (T1), 378 participants (204 women; mean age = 21.68, SD = 3.84) completed a 7-day smartphone-based ecological momentary assessment and questionnaires. A subsample (N = 216, 103 women; mean age = 21.79, SD = 3.74) additionally underwent fMRI. At follow-up (T2; 13.25 ± 8.37 months), 219 participants (112 women; mean age = 22.43, SD = 3.81) completed the same assessment protocol (164 with available fMRI data). Further demographic details are shown in Table 1, the Supplementary Information and Table S1. Participants had no history of neurological, general medical, or psychiatric disorders, as determined by clinical interviews^28^ and did not report any psychiatric symptoms. All subjects provided written informed consent for a protocol that was approved by the Heidelberg University institutional review board.

### Psychological assessment

Substance use patterns and contexts were assessed retrospectively using the ESPAD (European School Survey Project on Alcohol and Other Drugs)^31^ questionnaire. Current alcohol consumption was operationalized as standard drinks per week, and was included as a predictor in all models. This measure captures consumption quantity but does not assess alcohol-related impairment, loss of control, craving, tolerance, withdrawal, or consequences. Thus, the outcome should not be equated with alcohol-related problems or AUD severity. Only participants who self-identified as current drinkers were included in the analyses. Additional substance-use variables included *individual polysubstance use*, calculated as the sum of coded substance-use frequencies over the preceding 12 months, and a *risky substance-use environment*, reflecting the number of different substances used by close friends. Additional psychological measures are summarized in Table 1 and detailed in the Supplementary Information and Table S1.

### Ecological momentary assessments (EMA)

We assessed real-life social interactions and momentary affective valence using validated EMA scales^7, 49^ delivered via study smartphones (Motorola Moto G) across 7 consecutive days, with 5–17 possible prompts per day, accounting for a school-adapted sampling schedule for adolescents (see Supplementary Methods, Section 2.2; Fig. 2A). Momentary valence was captured as the average of two bipolar ratings (content– discontent; unwell–well). For social context assessment, participants reported whether they were alone or with others^7^. Age-adapted response scales were harmonized across participants. See Supplementary Methods for further details.

### Neuroimaging

We used a well-established incentive delay task during fMRI, which reliably evokes VS network responses^29^. The task consisted of monetary (MID) and social reward (SID) components and two task conditions (cue win and cue neutral, see Figure S1). We did not differentiate between reward types based on previous evidence of overlapping neural substrates^29^. Functional images were acquired using an echo-planar imaging sequence on a 3 Tesla scanner (Siemens Trio) and preprocessed using standard routines in SPM12 (http://www.fil.ion.ucl.ac.uk/spm/). See Supplementary Methods for further details.

### Data analysis and statistical inference

We analyzed questionnaire data using Spearman correlations and multiple linear regression models (IBM, SPSS, version 25).

To test whether the daily affective benefit from social interaction (vs. alone time) varied across individuals’ alcohol consumption levels—as operationalized by the alcohol × social context interaction—we examined main and interaction effects of momentary social contact (alone vs. in company) and alcohol consumption (ESPAD questionnaire; see Section 2.1) on momentary affective valence using multilevel models^7^. Models included time of day and time of day squared as covariates and were adjusted for compliance. We conducted all analyses in SAS 9.4 (SAS Institute), with an alpha level of 0.05.

We processed fMRI data in SPM12 by combining SID and MID runs into one general linear model (GLM) at the single-subject level. We modeled reward anticipation (win cues vs. neutral cues) to compute beta images at the first level. We included alcohol consumption quantity as a predictor in second-level multiple regression models. Statistical significance was set at p < 0.05, peak-level FWE-corrected within the predefined VS mask^29^. We examined task-dependent connectivity of the VS using psychophysiological interaction (PPI) analysis within the same VS mask. PPI interaction terms (physiological × psychological) for each task condition were entered into a GLM with task regressors, movement regressors and session constants. The PPI win cue > PPI neutral cue contrast was tested at the first level and entered into second-level random effects regression with alcohol consumption quantity as predictor of interest.

Due to the skewed distribution of the main predictor (alcohol consumption), we applied log transformation, non-parametric statistics, or residual checks where relevant (Fig. S2). To quantify changes in actual drinking amounts from baseline to follow-up, we used untransformed values (standard drinks/week).To account for age-related variation in alcohol use, age was included as a covariate in all primary models. We also ran an additional sensitivity analysis testing for age-by-alcohol interactions to ensure that the findings are not driven by age-specific effects (Supplementary methods). For transparency and comparability, subgroup comparison of key behavioral and imaging parameters are reported in the Supplement (Table S3). All models also accounted for sex, given known sex-specific effects of alcohol^50^. Control analyses included legal alcohol access (yes/no) as an additional covariate; findings were robust to this adjustment (Supplementary results).

To identify factors associated with change in alcohol consumption, we used change-score (Δ T2–T1) approach: we regressed changes in alcohol consumption on corresponding changes in measures, controlling for baseline alcohol use, age, and sex. This approach captures within-person temporal covariation identifying factors that vary in tandem with drinking behavior over time^48^. Control analyses confirmed result stability (Supplementary results and Table S4).

Where applicable, inferential p-values were adjusted for multiple testing using the Benjamini– Hochberg procedure within modality-specific hypothesis families. Descriptive analyses were exploratory and not used for inference (Table S4).

## Data availability

The data that support the findings of this study are available from OB upon reasonable request.

## Code availability

The custom code used for the analyses of this study is available from OB upon reasonable request.

## Acknowledgments

We thank Beate Höchemer, Mirjam Melzer, Urs Braun, Janina I. Schweiger, Sascha Moldavski and our research assistants for valuable support. We would also like to thank all of the participants and their families for their time and support in this study.

## Author contributions

HT, AML and UEP concepted and designed the study. OB, GG, MR recruited the participants and collected the data. OB, GG, AG, performed the data analyses. OB, AML, HT and FK drafted the manuscript. OB, GG, JA, MR, AG, UEP, CH, AML, HT and FK interpreted the data. All authors revised the manuscript critically for important intellectual content and approved the final version.

