## Supplement for "Multimodal longitudinal prediction of alcohol-use trajectories in adolescents and young adults: Social context, real-life social affective benefit, and fronto-striatal involvement"

[#]These authors contributed equally

### Corresponding Author

### Content


1. Participants and study design 2
2. Methods 2
    1. Socio-demographic and psychological inventory 2
    2. Ecological momentary assessments 2
        - Hardware and e-diary sampling 3
        - E-diary Measures 3
        - Multilevel modeling of e-diary assessments 3
    3. Neuroimaging 5
        - Reward anticipation task 5
        - fMRI data acquisition, preprocessing, and analysis 5
        - fMRI data analysis 5
        - fMRI data quality control 5
        - Mask 5
3. Supplementary Results 6
4. Supplementary Tables 7
5. Supplementary Figures 14
6. References 16

### 1. **Participants and study design**

Data for this analysis were drawn from the psychoepidemiological center (PEZ) at the Central Institute of Mental Health in Mannheim, Germany, a longitudinal multimodal community-based study. Participants, aged from 12 to 28, were recruited at baseline via the local population registry from local communities in the Rhine-Neckar metropolitan area, taking into account stratification of the population according to age, sex and nationality[1]. Data collection for the baseline assessment occurred between September 2014 and March 2019, and follow-up assessments were conducted between November 2015 and December 2020. The mean inter-assessment interval was 13.25 ± 8.37 months (minimum/maximum = 7/60 months). A total of 219 participants completed both assessment waves.

General exclusion criteria at baseline were the presence of a significant general medical or neurological diseases, drug or alcohol abuse, head trauma, and reported standard magnetic resonance imaging (MRI) contraindications (e.g., metal implants and pregnancy). Participants were further excluded from data analysis if they endorsed a current or lifetime psychiatric disorder (e.g., major depression, anxiety disorder, schizophrenia) as determined by the Mini-DIPS or the SCID-IV interview [2-4]. In the final analysis we excluded participants (both, baseline between-subject and the within-subject longitudinal analyses) if the following data quality control criteria applied: (i) e-diary compliance below 30%, or (ii) insufficient quality MRI data (see details below). Participants with missing questionnaire datasets were excluded from the analysis. This resulted in the sample (N = 378, 204 women; mean age = 21.68, SD = 3.84) for baseline and (N = 219, 112 women; mean age = 22.43, SD = 3.81) for follow-up (For details see Table 1). All participants provided informed consent and received monetary compensation.

Each participant admitted to the study completed an identical assessment protocol at each wave, consisting of an online questionnaire battery followed by a 7-day EMA sampling in everyday life. The MRI session took place on average 1.56 ± 2.10 days (minimum/maximum = 0/8 days) after the end of the EMA study week.

### 2. **Methods**

#### 2.1. **Socio-demographic, psychological and substance use measures**

An online questionnaire battery included basic socio-demographic measures and social-psychological measures including among others aggregated index of socioeconomic status (SES [5]), social network size (SNI)[6]. The detailed overview of the self-reported battery used in the study given in the Table S1.

Substance-use measures were derived from items used in the European School Survey Project on Alcohol and Other Drugs (ESPAD)[7]. Established in the 1990s, ESPAD is a large-scale European study designed to collect data on substance use among adolescents and young adults across Europe. The standard ESPAD battery comprises questions covering alcohol, tobacco, and illicit drug consumption (approx. 100 substance-specific items)[7]. *Alcohol consumption* was quantified as a continuous summary score reflecting self-reported standard drinks per week. Responses for four beverage categories (beer, wine, spirits, and cocktails) were converted from the original response categories (0–6) into estimated drinks per week and summed to obtain total weekly consumption. *Risky substance-use environment* was operationalized as a composite score indexing the prevalence of substance use among close friends. *Polysubstance use* captured participants' own substance use and was calculated as the sum of coded frequency responses for each assessed psychoactive substance over the preceding 12 months, with non-use coded as zero. The fictitious substance Relevin, included as a control item for potential over-reporting, was excluded from the polysubstance-use score. The complete item wording in the original German version and corresponding English translations is provided in Table S2.

Variables for cross-sectional and longitudinal characterization were selected based on their established relevance to alcohol use and change over time, and the phenotypic scope of the cohort[8-11].

We analyzed questionnaire data using Spearman correlations and multiple linear regressions (IBM, SPSS, version 25). Since the distribution of alcohol consumption—a central variable in our analyses—was skewed (see Figure S2), we applied a log10 transformation with an offset to accommodate zero values, and, where appropriate, used non-parametric statistics or evaluated model residuals. All models included sex and age as covariates of no interest.

#### 2.2. **Ecological momentary assessments**

*Hardware and e-diary sampling.* Participants carried a study smartphone (Motorola Moto G; Motorola Mobility) with electronic diary assessments and wore a hip-mounted accelerometer (Move-II or Move-III; movisens GmbH) for seven consecutive days in daily life. Before the start of the assessment period, participants were instructed and trained in the use of all devices.
We used a combined time- and location-contingent prompting protocol aimed at improving spatial coverage and increasing within-person variability of assessments[1]. Location-contingent prompts were generated by a smartphone algorithm that continuously tracked successive positions and issued a prompt once displacement from the previous location exceeded 500 m. In addition, we scheduled two fixed daily prompts (08:00 and 22:20). Prompts were signaled acoustically and visually and by vibration, and participants could delay responding by 5–15 min. We implemented the e-diaries in movisensXS (version 0.6.3658; movisens GmbH; https://xs.movisens.com).
Across study days, prompting occurred between 07:30 and 22:30, yielding 5–17 prompts per day with a minimum inter-prompt interval of 40 min and a maximum interval of 100 min[1,12,13]. For participants attending school, prompting was scheduled only for time periods outside regular school hours.
*E-diary Measures.* In adults, we assessed affective valence in daily life using a validated, widely used two-item EMA scale with good reliability and sensitivity[14]. The scale included two bipolar items ("content"–"discontent" and "well"–"unwell"), presented as computerized visual analogue scales ranging from 0 to 100, with reversed polarity at the anchors [14]. We instructed participants to place, when e-diary prompts were triggered, the locators at the scale positions representing their momentary affective state. We averaged the two items for each score to use as dependent variable in our multilevel analyses.
In adolescents, we assessed momentary affective valence using a 20-item scale sensitive to affective fluctuations and comprehensible for children [15]. Valence was measured with nine items on two affective dimensions 'good mood' (happy, satisfied, delighted, great, and cheerful) and "bad mood" (unhappy, angry, bad, afraid) [15]. Ratings were provided on 8-point Likert scales from 0 (not at all) to 7 (very intense). We computed subscale scores as the mean of corresponding items and rescaled them to a 0–100 range for comparability with adult measures. In addition, we assessed momentary social contact at each prompt[13,16]. Participants indicated whether they were alone or in the company of others at the moment of the e-diary rating (dichotomous measure). To ensure comparability, adolescent responses were rescaled to a 0–100 range.

*Multilevel modeling of e-diary assessments.* To examine whether alcohol consumption moderates the established association between momentary social contact (alone vs. in company) and affective valence—referred to as the social affective benefit[13] (SAB) — we estimated a random-intercept multilevel model in SAS (version 9.4; SAS Institute Inc., Cary, NC, USA), nesting repeated e-diary assessments (Level 1) within participants (Level 2). The model included alcohol consumption, dichotomous social contact [alone/in company], and their interaction as fixed effects. Time of day (linear and quadratic items, transformed to the daily study start time at 7:30 AM) were included as a Level-1 covariate[1,17]. Age, sex, compliance, and EMA protocol (adolescent vs. adult, to account for methodological differences in EMA assessments) were modeled at Level 2. The full model specification is shown in Equation S1.

*Equation 1 (main model):*

$$Y(\boldsymbol{affective\ valence})_{ij} = \beta_{00} + \beta_{01}{*}\text{Alcohol consumption}_j + \beta_{02}{*}age_j + \beta_{03}{*}sex_j + \beta_{04}{*}compliance_j$$
$$+\beta_{05}{*}protocol_j + \beta_{10}{*}social\ contact_{ij} + \beta_{20}{*}time\ of\ day_{ij} + \beta_{30}{*}time\ of\ day^2_{ij}$$
$$+ \boldsymbol{\beta_{40}{*}\textbf{Alcohol consumption}_j{*}social\ contact_{ij}} + u_{0j} + r_{ij}$$

In this model, $Y_{ij}$ denotes the affective valence reported by person *j* at time point *i*. Observations at each prompt (subscript *i*) are nested within individuals (subscript *j*). Beta coefficients represent fixed effects, including the intercept, the effect of our main predictor's alcohol consumption (assessed retrospectively using the ESPAD questionnaire; see Section 2.1), social contact (alone/not-alone), and their interaction, as well as the effects of the level-1 covariates time of day (linear and quadratic terms). Between-subject variables (Level 2) include age, sex, compliance and sampling protocol. The residual term $r_{ij}$ captures within-subject error. Random effect, i.e., individual-level deviations from the sample mean, is represented by $u_{0j}$ for the intercept.
Inspection of Level-1 residuals indicated no substantial deviation from normality (Supplementary

Fig. S3), suggesting that the model adequately captures the data structure in this sample.

#### 2.3. Neuroimaging

*Reward anticipation task.*

During fMRI, participants performed an adapted event-related reward anticipation task[18] assessing responses to monetary[19,20] and social[21] incentives (see Figure S1). Each trial began with a cue indicating either a potential reward or a neutral outcome, followed by a brief visual target to which participants responded as quickly as possible via button press. Upward-pointing arrows signaled reward availability, whereas sideways arrows marked neutral trials. On reward trials, responses delivered within a predefined time window triggered positive feedback, whereas neutral trials offered no reward. To equalize reward exposure across participants, the task dynamically adjusted the response deadline throughout the experiment, maintaining an average success rate of approximately 60%. The task provided feedback in the form of a €2 during the monetary reward task (MID) and a smiling female face during the social reward task (SID); blurred control images followed neutral trials and unsuccessful reward trials. The task comprised 30 trials in total (15 reward, 15 neutral) presented in pseudorandom order in each session. Participants completed social incentive delay and monetary incentive delay tasks as separate runs, with the SID always preceding the MID to minimize potential reductions in motivation after monetary reward exposure.

*fMRI data acquisition, preprocessing, and analysis.*

We acquired functional MRI (fMRI) data at CIMH on a 3 Tesla MRI scanner (Siemens Trio, Erlangen, Germany) with an echo-planar imaging (EPI) sequence (TE: 30 ms, TR: 2 s, flip angle: 80°, matrix: 64 × 64, FOV: 192 × 192 mm, in-plane resolution: 3 x 3 mm, slice thickness: 4 mm, 28 axial slices, 151 volumes). For both tasks we employed standard preprocessing routines implemented in SPM12 (http://www.fil.ion.ucl.ac.uk/spm), in which the images were realigned to the mean image of the scan run using a 6-parameter rigid body spatial transformation, spatially normalized to the standard stereotactic space of the Montreal Neurological Institute (MNI) template, resampled to 3 mm isotropic voxels and smoothed with an 8 mm full-width at half-maximum (FWHM) Gaussian kernel.

*fMRI data analysis*.

*Activation analysis*: We examined reward anticipation–related neural activity at the single-subject level by modeling differential responses to reward cues relative to neutral cues. The first-level general linear model (GLM) included two sessions - the social and monetary incentive tasks. For each session, we specified four task regressors corresponding to win cues, neutral cues, target presentation, and feedback. All task regressors were defined as stick functions and convolved with the canonical hemodynamic response function HRF. We included six head-motion parameters per session, derived from the realignment procedure, as nuisance regressors. During model estimation, we applied a high-pass filter with a cutoff of 128 s and accounted for temporal autocorrelations using a first-order autoregressive model. We computed contrast images comparing responses to win cues with responses to neutral cues (win cue > neutral cue). At the group level, we entered these contrast images into a multiple regression analysis to examine associations between reward anticipation–related activation and alcohol consumption while including age and sex as covariates of no interest.

*Psychophysiological interaction*. We used psychophysiological interaction (PPI) analysis to examine task-dependent connectivity[22] of the bilateral ventral striatum (VS) during the same reward anticipation task with two sessions as described above. We used the same mask (p.6.Mask) to extract the physiological regressor: the first eigenvariate of the BOLD signal residuals from voxels within this mask, obtained after the initial activation GLM. PPI interaction regressors were generated for each task condition separately (bilateral VS physiological timeseries convolved with win cue, neutral cue, flash/target, and feedback onsets). A PPI model was specified using the bilateral VS seed. It included the four PPI interaction regressors, the VS physiological timeseries (VOI), alongside the original four task regressors from the activation GLM, six movement parameters, and session-specific constants (total: 17 regressors). The PPI win cue > PPI neutral cue interaction contrast was tested at the first level. For group inference, the contrast images (PPI win cue > PPI neutral cue) were entered into a linear regression analysis in which we defined alcohol consumption as predictor of interest and included age, sex as covariates of no interest.

*fMRI data quality control.*

To ensure high data quality, we applied a standardized neuroimaging quality control pipeline. Head motion was quantified based on established metrics including the maximum translational and rotational displacement between volumes, as well as mean voxel-wise framewise displacement (FD)[18,23,24]. Participants were excluded if mean FD exceeded 0.5 mm or if more than 20% of volumes exceeded this threshold at either assessment time point (T1 or T2), following previously reported criteria. Additionally, signal quality was evaluated using the NYU Center for Brain Imaging Quality Assessment Toolbox, which provides signal-to-noise ratio (SNR)[25].

*Mask.*

In line with our hypothesis, significance for the incentive delay task was defined as $p < 0.05$, family-wise error (FWE) corrected within a well-established a priori defined mask of the VS[23]. The mask was defined by combining two anatomical regions: the caudate head from the WFU PickAtlas (human-atlas TD Brodmann areas) and the "accumbens" mask from the Harvard–Oxford Subcortical Structural Atlas[26]. To improve reliability, it was further constrained based on voxel-wise test–retest data from an independent sample (n = 28; mean age: 22.9 ± 2.8 years; 14 females)[27]. Only voxels with an intraclass correlation coefficient (ICC) above 0.4 were retained. Outside this ROI, whole-brain results were considered significant if they survived FWE correction at $p < 0.05$.

#### 2.4. Longitudinal analysis.

To identify factors associated with change in alcohol consumption, we modeled within-person change over time as a function of corresponding change in determinants examined in this study. Specifically, we used a change-score regression approach, regressing the change in alcohol consumption (Δ T2–T1) on the change in measures (Δ T2–T1) while controlling for baseline alcohol use, age, and sex. This model captures intra-individual co-variation between changes in individual-level factors and concurrent shifts in drinking behavior, emphasizing dynamic processes that evolve within the same individuals rather than stable inter-individual differences.

For momentary social affective benefit (SAB), we fitted separate multilevel models at T1 and T2, predicting momentary affective valence from social contact (alone vs. in company), time of day, and time of day squared (Level 1), adjusting for age, sex, EMA compliance, and protocol (Level 2). The models included participant-specific random intercepts and random slopes for social contact, from which individual SAB estimates were derived. Individual SAB estimates were then used to calculate within-person changes (T2–T1). For other EMA variables, we computed person-level summary statistics during each measurement week (T1, T2): means for continuous measures (e.g. affective valence) and proportion of EMA prompts for binary variables (social contact: time 'not alone' during each measurement week). For neural measures, the identical first-level GLM models for both activation (win cue > neutral cue) and PPI connectivity analyses (PPI win cue > PPI neutral cue) as described above were separately conducted for T2 data using the same preprocessing, task regressors, and VS ROI. We then extracted subject-specific VS activation betas (based on the T1 group-level peak coordinates showing an association between alcohol consumption and VS activation; ±6 mm spheres) consistently at T1 and T2. Following the same procedure, VS-dlPFC PPI parameter estimates (betas) were extracted (based on the T1 whole-brain PPI dlPFC peak coordinates showing an association with alcohol consumption (±6 mm spheres)) consistently from both timepoints. Then we calculated within-subject change (T2-T1) as described in Equation 2. The full model specification is shown in Equation 3.

*Equation 2:*

$$\text{Variable } \Delta = \text{Variable}_{T2} - Variable_{T1}$$

*Equation 3:*

$$\Delta \mathrm{Y} = \beta 0 + \boldsymbol{\beta} 1(\Delta \mathbf{X}) + \beta 2 \mathrm{Y}_{\mathrm{T1}} + \beta 3 \mathrm{age}_{\mathrm{T1}} + \beta 4 \mathrm{sex} + \varepsilon$$

In this model, Y- outcome variable (Alcohol use), $Y_{T1}$ - alcohol use measured at baseline (Time 1), ΔY - change in alcohol use between T1 and T2, calculated as shown in Equation 2, X – predictor (independent variable), ΔX - change in the predictor between T1 and T2, calculated as in

Equation 2, $Age_{T1}$ - age at baseline and Sex. Beta coefficients represent intercept, and regression coefficients for change in the predictor ($\Delta X$), for baseline alcohol use ($Y_{T1}$), and covariates, $\varepsilon$ - error term.

To address known limitations of difference-score models, including regression-to-the-mean artifacts and potential multicollinearity, all primary models included baseline alcohol use ($Y_{T1}$) as a covariate[28,29]. We further conducted a series of complementary and sensitivity analyses and reported descriptive statistics (means, SDs, T1–T2 correlations, Table S4.) to enhance interpretability and confirm result stability.

As a complementary analysis following the longitudinal analyses, we assessed the test–retest reliability of VS activation and VS–dlPFC connectivity across T1 and T2 using ICC(2,1) for absolute agreement and ICC(3,1) for consistency[23].

#### 2.5. Statistics

To account for age-related variation in alcohol use, we included age in all models. All models also accounted for sex, given known sex-specific effects of alcohol and the observed association between sex and alcohol use in our sample (Table 1). Legal access to alcohol was included in control analyses as a covariate to account for potential confounding due to differences in statutory drinking eligibility.
Where applicable, inferential p-values were adjusted for multiple testing using the Benjamini–Hochberg procedure[30] within modality-specific hypothesis families, and statistical significance was determined based on adjusted p-values. Descriptive analyses were used for exploratory characterisation of the data and were not used for inferential interpretation. Uncorrected and Benjamini–Hochberg–adjusted p-values for inferential models are provided in Table S4.

### 3. Supplementary Results

#### 3.1. Age effects

To ensure comparability across age groups, we examined whether sub-samples of adolescents (<18 years) and young adults (≥18 years) differed systematically on variables that could bias the brain–behavior-alcohol relationship—including psychological, behavioral, and neuroimaging metrics — beyond those that are inherently age-dependent or define the groups. To examine potential effects of differences in adolescent and adult EMA valence scales, we conducted a sensitivity analysis testing the interaction between ΔSAB and EMA protocol (adolescent vs. adult), which was not significant ($F(1, 148) = 1.50$, $p = .223$).

_Behavioral measures._ Comparisons between adolescent and young adult sub-samples revealed no widespread differences across key demographic or psychosocial measures, supporting overall comparability between these age-groups (all ps>0.100). Observed differences were limited to variables likely reflecting context-related variation rather than systematic age effects (Table S3 for more details).

_Task performance._ Mean reaction time analysis revealed a significant main effect of condition ($F(1,213) = 243.01$, $p < .001$), with faster responses following win compared to neutral cues. A main effect of age group was also observed ($F(1,213) = 7.75$, $p = .006$), with overall slower responses in adolescents, as commonly reported[31]. However, the condition × age group interaction was not significant ($F(1,213) = 0.04$, $p = .850$), suggesting that reward anticipation produced comparable behavioral effects across sub-samples (Table S3).

_Image quality metrics_. fMRI quality metrics did not differ between adolescent and adult sub-samples (all ps>0.070) confirming consistent data quality across groups (Table S3).

_Pre-scan affective state measures_. Pre-scan SAM ratings of valence, arousal, and dominance did not differ between adolescent and adult sub-samples (all ps > .42), further supporting comparable affective states prior to scanning (Table S3).

*Daily measure.* To rule out age-specific effects, we estimated a separate multilevel model testing whether age moderated the alcohol-social context interaction on momentary valence (three-way interaction: age × alcohol consumption × social context). This analysis revealed no significant moderation ($p = 0.225$), indicating no evidence of age-dependent moderation of alcohol-related affective benefits in social contexts.
*Reward-related fMRI responses*. We tested for age effects by modeling the interaction between alcohol consumption and age on VS activation and task-based connectivity. These analyses revealed no significant age-related modulation of neural responses and no evidence for age-dependent effects of alcohol consumption on activation or connectivity within the reward network (all ps > 0.63).

#### 3.2. Sex effects

To exclude potential sex-specific influences, we tested whether sex moderated the association between alcohol consumption, daily and neural measures. These analyses revealed no significant interaction effects, indicating that alcohol-related effects observed in daily and/or neural measures were not affected by sex (all ps > 0.179).

#### 3.3. Temporal variability of study measures.

To aid interpretation of change-score effects, we report descriptive statistics of T1–T2 changes (means, standard deviations, and correlations) for alcohol use and psychosocial predictors. As shown in Table S4, descriptive statistics indicated substantial variability in T1-T2 change across predictors (range of mean Δ values: -0.72 to +0.30; SDs 0.24-20.87), indicating sufficient variability in longitudinal change to enable the detection of individual differences. Temporal stability showed a clear gradient ($r/\varphi = .19\text{-}.94$), with psychosocial risk factors (risky environment $r = .39$; polysubstance $r = .19$) exhibiting low stability versus demographic markers (urbanicity $r = .94$). Taken together, these patterns—substantial Δ variability across factors and differing levels of temporal stability— support the use of change-score models to emphasize within-person processes rather than stable between-person differences. These descriptives thus support the interpretability of our Δ-Δ approach.
Test–retest reliability was fair-to-good for both VS activation (ICC(2,1) = .50, 95% CI [.36, .61]; ICC(3,1) = .51, 95% CI [.37, .62]) and VS–dlPFC connectivity (ICC(2,1) = .43, 95% CI [.26, .57]; ICC(3,1) = .46, 95% CI [.32, .58])[23].

### Table S1: Psychosocial assessments

| Variable | Description/Inventory | Scale |
|---|---|---|
| Socioeconomic status (SES) | Health Update 2009 (GEDA)[5]<br>Socioeconomic status assessment of the German | Multidimensional aggregated index of status-constituting dimensions (education, occupation, income) |
| Current urbanicity | Current place of residence[32] | Ordinal (1 = rural, 2 = town, 3 = city) |
| Urban upbringing | Early-life urbanicity (birth–15 years)[32] | Cumulative years-weighted urbanicity score (birth–15 years) |
| Daily stress | Alltagsbelastungsfragebogen (ABF)[33] | 58 items, 8-point Likert scales; daily sum scores averaged across 7 days. |
| Social network | Social network index[6] | Social network size (total number of individuals with regular social contact across up to 12 [illegible] |
| Body-mass-index [illegible] | calculated based on participants self-reported height and weight | |
| Well-Being (who) | WHO-5 Well-Being Index[34]<br>Perceived current mental well-being | 5 items, 6-point Likert scales |
| Satisfaction with Life | Satisfaction with Life Scale (SWLS)[35]<br>Global cognitive judgments of satisfaction with life | 5 items, 7-point Likert scales |
| Sense of Coherence | Sense of Coherence Scale (SOC-29)[36]<br>Dispositional sense of coherence | 29 items, 7-point Likert scales |
| Self-Efficacy | General Self-Efficacy Scale (GSES)[37]<br>Generalized belief in the own ability to control environmental challenges | 10 items, 4-point Likert scales |
| Perceived social [illegible] | Berlin Social Support Scale (BSSS)[38] | 8 items, 4-point Likert scales |

**Table S2. Substance Use Measures**

| Measure | Question (German / English) | Response options | Response format |
|---|---|---|---|
| Alcohol consumption (standard drinks per week) | Wie viel von diesen alkoholischen Getränken trinken Sie derzeit?<br>How much of the following alcoholic beverages are you currently drinking? | (a) *Bier* (beer);<br>(b) *Wein, Obstwe in, Sekt* (wine, fruit wine, sparkling wine);<br>(c) *Schnaps* (spirits);<br>(d) *Cocktails, Longdrinks* (cocktails, long drinks) | 7 response categories ranging from „Gar nicht“ (not at all) to „1 oder mehr Gläser tgl.“ (≥1 glass daily) |
| Risky substance-use environment | Wie viele Ihrer Freunde schätzen Sie…<br>How many of your friends do you estimate… | *(i) rauchen Zigaretten* (smoke cigarettes);<br>*(ii) trinken alkoholische Getränke (Bier, Cider, Alkopops, Wein, Spirituosen)* (drink alcoholic beverages);<br>*(iii) betrinken sich* (get drunk);<br>*(iv) rauchen Marihuana oder Haschisch (Cannabis)* (smoke marijuana or hashish);<br>*(v) nehmen Beruhigungs- und Schlafmittel (ohne ärztliche Anordnung)* (take sedatives and sleeping pills without prescription);<br>*(vi) nehmen Aufputschmittel (Ecstasy)* (take stimulants);<br>*(vii) schnüffeln Leim oder Lösungsmittel* (sniff glue or solvents) | 5 ordinal response categories reflecting proportion of peers (keiner, einige wenige, etwa die Hälfte, die meisten, alle) |
| Polysubstance use (self-use) | Haben Sie folgende Mittel in den letzten 12 Monaten ausprobiert und wenn ja, wie oft? Falls Sie ein Mittel in den letzten 12 Monaten nicht probiert haben, geben Sie bitte die „0“ (Null) an.<br>Have you tried the following substances during the past 12 months, and if so, how often? If you have not tried a substance during the past 12 months, please indicate “0” (zero). | *(i) Marihuana* (marijuana);<br>*(ii) Aufputschmittel (Amphetamine, Speed, MDMA, Ecstasy)* (stimulants);<br>*(iii) Medikamente, um sich zu berauschen* (medications used for intoxication);<br>*(iv) Schnüffeln von Leim oder Lösungsmitteln* (sniffing glue or solvents);<br>*(v) Beruhigungs- und Schlafmittel (ohne ärztliche Anordnung)* (sedatives and sleeping pills without prescription);<br>*(vi) LSD oder andere halluzinogene Substanzen* (LSD or other hallucinogens);<br>(vii) *Crack* (crack cocaine);<br>*(viii) Kokain* (cocaine);<br>*(ix) Relevin* (Relevin);<br>*(x) Heroin* (heroin);<br>*(xi) „Magic Mushrooms“/Pilze* (psilocybin-containing mushrooms);<br>*(xii) GHB / Liquid Ecstasy* (GHB / liquid ecstasy);<br>*(xiii) Anabole Steroide* (anabolic steroids) | Ordinal frequency scale per substance (0 to ≥40 times in the past 12 months) |

### Table S3: Comparison between age groups, N=378.

| Characteristics | Adolescents (n=80) (<18 years) | | | Young adults (n=298) (≥18 years) | | | t-test / MW U-test / $\chi^2$ / Fisher[a] |
|---|---|---|---|---|---|---|---|
| | M ± SD / count / % | min/max | n | M ± SD / count | min/max | n | *p* value[b] |
| ***Demographic data*** | | | | | | | |
| Sex (male/female) | 43/37 | N/A | 80 | 131/167 | N/A | 298 | 0.163 |
| Nationality (German/other) | 80/0 | N/A | 80 | 266/32 | N/A | 298 | **<0.001** |
| Current urbanicity | 2.18±0.82 | 1/3 | 80 | 2.46±0.70 | 1/3 | 297 | **0.002** |
| Urban upbringing | 32.00±11.84 | 13.00/45.00 | 80 | 29.72±11.02 | 14.00/45.00 | 295 | 0.109 |
| Household income (€/month after taxes) | 4281.25 ±1375.49 | 2125.00/6000.00 | 69 | 2509.47±1752.08 | 500.00/6000.00 | 293 | **<0.001** |
| Socioeconomic status (SES) | 16.41±2.93 | 8.70/21.00 | 80 | 14.43±3.24 | 5.5/21.00 | 298 | **<0.001** |
| Familial psychiatric history (yes/no) | 9/71 | N/A | 80 | 97/199 | N/A | 296 | **<0.001** |
| Handedness (left/right)[#] | 11/51 | N/A | 62 | 28/196 | N/A | 224 | 0.294 |
| ***Psychosocial Well-being and Stress*** | | | | | | | |
| Wellbeing (WHO-5, sum) | 15.44±3.75 | 5.00/23.00 | 79 | 15.92±4.19 | 5.00/25.00 | 291 | 0.258 |
| Satisfaction with life (SWLS) | 26.74±5.18 | 8.00/34.00 | 78 | 26.93±5.35 | 9.00/35.00 | 296 | 0.770 |
| Self-efficacy (SWE, sum) | 29.99±4.48 | 15.00/40.00 | 79 | 30.05±4.20 | 17.00/40.00 | 296 | 0.907 |
| Perceived social support (BSSS, sum) | 29.60±3.75 | 10.00/32.00 | 79 | 30.09±3.15 | 8.00/32.00 | 297 | 0.246 |
| Sense of coherence (SOC, sum) | 139.28±22.55 | 76.00/182.00 | 80 | 146.82±20.78 | 64.00/190.00 | 296 | **0.005** |
| Daily Stress (ABF, mean daily sum) | 33.00±25.13 | 2.00/125.71 | 71 | 28.57±20.71 | 0.43/159.29 | 256 | 0.131 |
| Social network size (SNI, individuals )[#] | 16.67±5.94 | 7/36.00 | 49 | 15.60±5.85 | 0/37.00 | 183 | 0.264 |
| ***EMA*** | | | | | | | |
| Social contact (not alone, %)[c] | 56.21 | N/A | 80 | 52.79 | N/A | 298 | **<0.001** |
| E-diary prompts across 7-days | 36.39±9.58 | 14.00/73.00 | 80 | 68.62±17.38 | 18.00/170.00 | 298 | **<0.001** |
| E-diary Compliance (%)[d] | 80.03±15.09 | 37.84/100 | 80 | 79.97±16.56 | 30.51/100 | 298 | 0.978 |
| ***fMRI quality parameters*** | | | | | | | |
| Number of available scans | N/A | N/A | 44 | N/A | N/A | 172 | N/A |
| Signal to noise ratio | 88.19±14.84 | 56.00/110.72 | 44 | 84.37±16.68 | 45.90/125.43 | 172 | 0.172 |
| Movement: total translation (mm) | 0.75±0.46 | 0.19/2.53 | 44 | 0.63±0.35 | 0.15/2.53 | 172 | 0.070 |
| Movement: total rotation (degree) | 0.64±0.38 | 0.19/2.05 | 44 | 0.58±0.44 | 0.17/3.39 | 172 | 0.385 |
| Mean FWD (mm) | 0.17±0.07 | 0.08/0.34 | 44 | 0.16±0.06 | 0.07/0.32 | 172 | 0.248 |
| ***fMRI task performance*** | | | | | | | |
| Reaction time (ms) | 257.73±26.58 | 217.07/338.91 | 44 | 246.74±20.11 | 201.00/332.21 | 172 | **0.003** |
| Number of win trials (max 15) | 10.33±1.85 | 7.50/12.00 | 44 | 10.22±1.4 | 7.00/13.50 | 172 | 0.588 |

| | | | | | | | |
|---|---|---|---|---|---|---|---|
| ***Subjective affective state (pre-scan SAM)*** | | | | | | | |
| SAM Arousal rating | 3.75±0.86 | 2.00/5.00 | 44 | 3.81±0.75 | 2.00/5.00 | 172 | 0.625 |
| SAM Dominance rating | 3.81±0.73 | 3.00/5.00 | 44 | 3.72±0.66 | 2.00/5.00 | 172 | 0.423 |
| SAM valence rating | 1.89±0.65 | 1.00/3.00 | 44 | 1.84±0.55 | 1.00/3.00 | 172 | 0.614 |

For details on the psychological questionnaires including acronyms, see Supplemental Table S1. Abbreviations: M = mean, SD = standard deviation, n = number of participants with available data; N/A (Not applicable) → the value cannot logically exist.
[#] - available for a subsample of participants.
[a] t-test = t-test for independent samples, χ2 = chi-squared test, Fisher = Fisher's exact test (used when expected cell counts < 5). U-test = Mann-Whitney U-test. Group comparisons were conducted without covariate adjustment for descriptive purposes.
[b] *p*-values are uncorrected and reported for descriptive purposes.
[c] Percentage of completed EMA prompts spent in company across 7 days of measurement.
[d] Percentage of answered prompt.

### Table S4. Descriptive statistics and inferential results for longitudinal change analyses

| Measures | Longitudinal (N=219)[a] | | | | | | |
|---|---|---|---|---|---|---|---|
| | T1 | T2 | Mean ±SD Δ /status change | Stability (T1, T2) | change model (Δ–Δ) | | |
| | **Mean ± SD**/ count | **Mean ± SD**/ count | | r/φ[b] | p-value (uncorr.) | p-value (BH-FDR) | **95 % CI** (unadjusted) |
| Risky substance-use environment | 4.54±2.25 | 3.82±2.98 | -0.72 ±2.95 | 0.394 | **<0.001** | **0.004** | 0.046 – 0.067 |
| Individual polysubstance use | 1.46±2.49 | 1.67±5.79 | 0.20±5.85 | 0.192 | **<0.001** | **0.004** | 0.010 - 0.022 |
| Smoking (nonsmoker/smoker) | 137/29 | 133/33 | 146/20[c] | 0.605 | **0.013** | **0.026** | 0.031 – 0.298 |
| Daily stress (ABF, mean daily sum) | 30.41±23.88 | 29.72±25.37 | -0.70±20.87 | 0.642 | 0.131 | 0.150 | -0.006 – 0.007 |
| Social network size (SNI, individuals) | 16.04±5.78 | 16.35±5.96 | 0.30±5.48 | 0.565 | **0.005** | **0.013** | 0.003 – 0.019 |
| Perceived social support (BSSS, sum) | 29.99±3.29 | 29.80±4.05 | -0.22±4.10 | 0.387 | 0.087 | 0.116 | -0.016 – 0.001 |
| Socioeconomic status (SES) | 14.91±3.26 | 14.99±3.33 | 0.09±0.84 | 0.968 | N/A | N/A | N/A |
| Current urbanicity | 2.47±0.72 | 2.47±0.73 | 0.004±0.24 | 0.943 | **0.041** | 0.066 | 0.007 – 0.317 |
| Body mass index (BMI, kg $m^{-2}$) | 22.87±4.75 | 23.09±4.09 | 0.22±3.10 | 0.767 | 0.466 | 0.466 | -0.017 – 0.008 |
| Urban upbringing | --- | --- | --- | --- | N/A | N/A | N/A |
| Familial psychiatric history | --- | --- | --- | --- | N/A | N/A | N/A |

Abbreviations: SD – standard deviation, Δ – delta (T2-T1); CI - Confidence Interval.
[A] Descriptive statistics were calculated using available data for each measure.
[b] φ coefficient for dichotomous scales; Pearson r for continuous scales;
[c] For smoking, status change is reported as unchanged/changed.

### Table S5: Multilevel analysis results

| Dependent variable | Predictor | Estimate | Standard Error | DF | T value | 95 % CI | P value |
|---|---|---|---|---|---|---|---|
| **Valence** (0-100) | Intercept | 63.326 | 4.482 | 386 | 14.13 | 54.515 – 72.138 | <0.001 |
| | Time (hours) | 0.539 | 0.089 | 23E3 | 6.06 | 0.365 – 0.714 | <0.001 |
| | Time-squared (hours$^2$) | -0.017 | 0.006 | 23E3 | -2.96 | -0.028 – -0.005 | 0.003 |
| | Sex: | | | | | | |
| | male | 0.661 | 1.184 | 371 | 0.56 | -1.668 – 2.990 | 0.577 |
| | female | 0 | - | - | - | - | - |
| | Age (years) | -0.074 | 0.222 | 369 | -0.34 | -0.512 – 0.362 | 0.737 |
| | Compliance | 0.093 | 0.035 | 402 | 2.64 | 0.024 – 0.162 | 0.009 |
| | Measurement protocol | -1.368 | 2.097 | 374 | -0.65 | -5.491 – 2.755 | 0.515 |
| | Social contact | | | | | | |
| | In company | 2.445 | 0.239 | 23E3 | 10.21 | 1.976 – 2.914 | <0.001 |
| | Alone (reference category) | 0 | - | - | - | | |
| | **Alcohol consumption** | **-3.117** | **1.356** | **399** | **-2.30** | **-5.784 – -0.451** | **0.022** |
| | **Alcohol consumption*social contact** | | | | | | |
| | **In company** | **1.274** | **0.476** | **23E3** | **2.68** | **0.341 – 2.206** | **0.007** |
| | **Alone (reference category)** | 0 | - | - | - | - | - |

Abbreviations: df = degree of freedom, CI = confidence interval. Beta coefficients (β) represent unstandardized fixed-effect estimates. All reported p values for beta coefficients are two-sided and derived from the t-statistics of the multilevel model.
The multilevel model included 23,393 EMA observations from 378 participants.

### Supplemental Figures

**Figure S1: Reward anticipation task.**

Subjects were asked to give speeded responses to a visual target (brief light flash). Preceding cues indicated whether subjects have the chance to win 2 Euros (monetary condition) or Julia's smile (social condition) if the response is fast enough. Note that reward types were presented in two separate sessions which were combined during data analysis. ITI: inter trial interval.

**Figure S2: Baseline Distribution of Alcohol consumption**

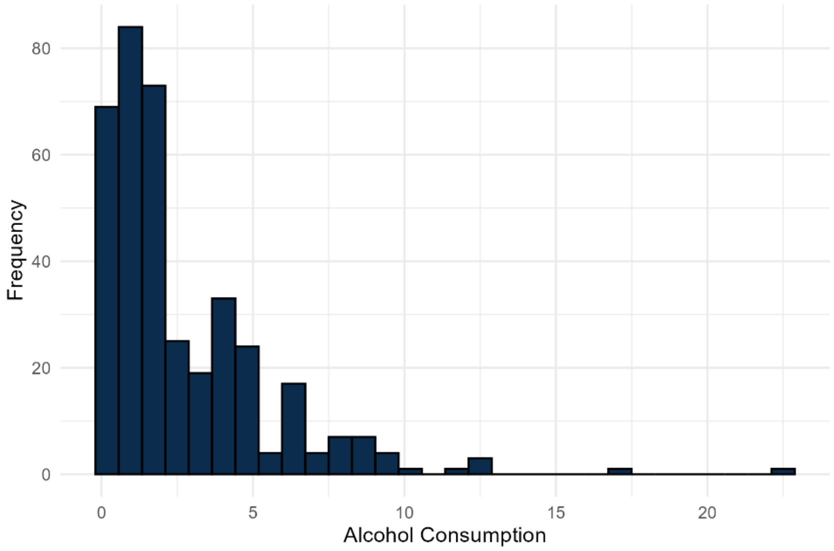


The histogram depicts the distribution (y-axis - frequency) of alcohol consumption (standard drinks/week, x-axis), indicating the skewed distribution of the measure.

**Figure S3: Distribution of level-1 residuals of the multi-level model**

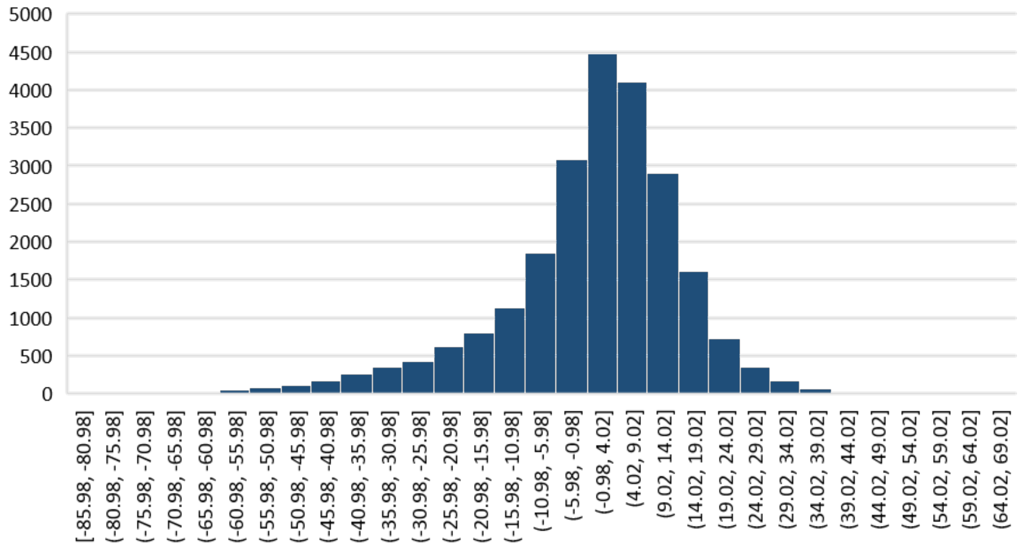


The histogram illustrates the distribution (y-axis shows the frequency) of level-1 residuals (x-axis), which measure deviations from the conditional mean (conditional residuals) derived from multi-level model (see Supplementary Methods: 2.2. Ecological momentary assessments, and Table S5). Visual inspection indicated no substantial deviation from normality, suggesting that the model adequately captures the data structure in this sample.

**Figure S4: Total number of prompts participants have been alone and not alone at any given e-diary prompt across 7 days of measurement.**

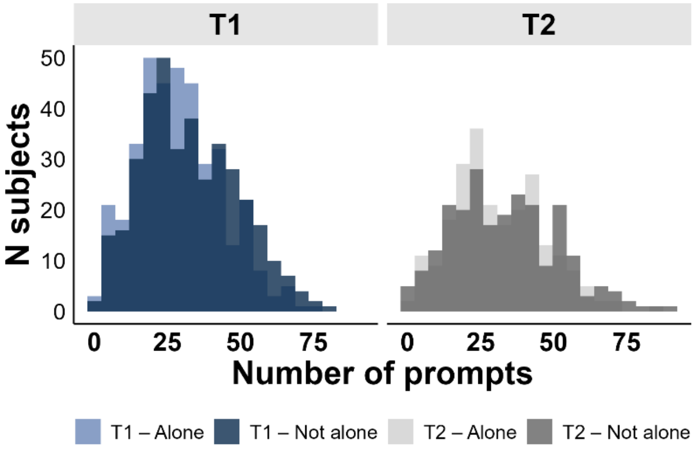


Distribution of the number of ecological momentary assessment prompts completed at baseline (T1) and follow-up (T2), shown separately for assessments conducted alone and not alone. Histograms display the frequency of completed prompts per participant, illustrating broadly similar distributions of completed prompts between T1 and T2 within each social context (alone vs. not alone).
